# Essentiality of *CREBBP* in *EP300* truncated B-cell lymphoma revealed by genome-wide CRISPR-Cas9 screen

**DOI:** 10.1101/746594

**Authors:** Man Nie, Likun Du, Bo Zhang, Weicheng Ren, Julia Joung, Xiaofei Ye, Jonathan Arias Fuenzalida, Xi Shi, Dongbing Liu, Kui Wu, Feng Zhang, Qiang Pan-Hammarström

## Abstract

Histone acetyltransferases (HATs), including *CREBBP* and *EP300*, are frequently mutated in B-cell malignancies and usually play a tumor-suppressive role. In this study, we performed whole genome and transcriptome sequencing and a genome-wide CRISPR-Cas9 knockout screen to study a germinal center B-cell like diffuse large B-cell lymphoma (DLBCL) cell line (RC-K8). Using a summarizing method that is optimized to address the complexity introduced by the time-course design, we identified a distinct pattern of genetic essentialities in RC-K8, including a dependency on *CREBBP* and *MDM2*, shown already at early time points and a gradually increased dependency on oxidative phosphorylation related genes. The dependency on *CREBBP* is associated with the corresponding genetic alterations identified in this cell line, i.e. a balanced translocation involves *EP300*, which resulted in a truncated form of protein that lacks the critical bromodomain and HAT domain. We further evaluated the previously published CRISPR-Cas9 screens and identified a genetic essentiality of *CREBBP* or *EP300* gene in a small set of cancer cell lines, including several DLBCL cell lines that are highly sensitive for *EP300* knockout and with *CREBBP* mutations or copy number loss. The dependency of the remaining HAT function in *CREBBP* and/or *EP300*-deficient genotype was validated by testing the HAT-domain inhibitor A-485. Our study suggests that integration of the unbiased, time-course-based functional screen results with the genomic and transcriptomic data can identify druggable vulnerability in individual or subgroups of cell lines/patients, which may help to develop more effective therapeutic strategies for cancers that are genetically highly heterogeneous, like DLBCL.

## Introduction

Diffuse large B-cell lymphoma (DLBCL) is one of the most common types of aggressive lymphoid malignancy. With the current standard immuno-chemotherapy, around 30-40% of DLBCL patients still suffer from refractory disease or relapse (1, 2). Based on transcriptional profiles, two major subtypes of DLBCL have been defined: GCB (germinal center B-cell like) and ABC (activated B-cell like) (1). Large-scale genome sequencing has further enabled identification of several molecular subtypes of DLBCL based on genetic alterations, affecting the proto-oncogenes *BCL-2/-6* and *MYC*, epigenetic modifiers and regulators in the B-cell receptor (BCR), NF-κB, NOTCH and p53 signaling pathways (3, 4). The ABC subtype and selected molecular subtypes (C3 and C5 in Chapuy *et al*. (3); MCD and N1 in Schmitz *et al*. (4)) are associated with a poor prognosis. Recently, we have shown that hepatitis B virus (HBV)-related DLBCLs are associated with unique genetic and clinical features as well as a shorter overall patient survival, and may be considered as a distinct subtype (5). DLBCL is thus a highly heterogeneous disease, and identification of genetic vulnerabilities that are specific to a subtype or subgroup of patients will aid the development of novel targeted therapeutic strategies and improve clinical outcome.

One recently developed promising approach to systematically identifying such genetic vulnerabilities is through genome-wide CRISPR-Cas9 screening (6), which has been applied in a range of contexts to discover genetic dependencies and vulnerabilities in cancer cells (7–9). In the context of B-cell malignancy, CRISPR-Cas9 screens have already provided valuable results for understanding the mechanisms of tumorigenesis (10, 11).

Dysregulation of epigenetic modulators is frequently observed in lymphomas. *EZH2*, *KMT2D*, *MEF2B*, as well as *CREBBP*, the histone acetyltransferase (HAT) encoding gene and its paralogue *EP300* are among the most frequently mutated genes in B-cell lymphomas (12–14). Despite the sequence homology and functional similarities between *CREBBP* and *EP300* (15), monoallelic germline lesions in either gene may cause a severe phenotype, i.e., Rubinstein-Taybi syndrome (16). In B-cell lymphomas, heterozygous somatic mutations in either gene may result in haploinsufficiency (17, 18). Genetic alterations in *CREBBP* are, however, more frequently observed in these disorders than deficiency of *EP300*. Recently, functional screening by a siRNA library suggested that *EP300* is a specific synthetic lethal gene in *CREBBP*-deficient lung cancer cells, raising the possibility to use *EP300* as a novel therapeutic target for treatment of *CREBBP*-mutated tumors (19).

In this study, we performed whole genome and transcriptome sequencing to characterize a GCB-like DLBCL cell line (RC-K8), established from peritoneal effusions of a patient with terminal, refractory stage of disease (20). We furthermore applied an unbiased, genome-wide CRISPR-Cas9 loss-of-function screening approach to study genetic dependencies in this cell line. For comparison, we have also evaluated the published CRISPR-Cas9 screens from a pan-cancer study and sets of DLBCL cell lines. We identified a distinct pattern of genetic essentialities in RC-K8, including a specific dependency on *CREBBP* in the context of *EP300* deficiency. We also observed essentiality of *EP300* in several DLBCL cell lines harbor mutation, translocation or copy number loss in *CREBBP*. Our results suggest that the dependency of the remaining HAT function is a druggable vulnerability in *CREBBP-* or *EP300-*deficient DLBCLs.

## Materials and Methods

### Cell culture

DLBCL cell lines RC-K8, OCILY3, SUDHL4, SUDHL10 and Pfeiffer were purchased from the Leibniz-Institute DSMZ (Braunschweig, Germany) or the American Type Culture Collection (ATCC, Manassas, USA). U2932 was kindly provided by Dr. G. Enblad’s research group (Uppsala University). All cell lines were cultured in RPMI 1640 (Invitrogen, Carlsbad, USA) supplemented with 10% FBS (Gibco, Invitrogen, Paisley, UK).

### Whole genome sequencing

DNA from RC-K8 cells was sequenced using the BGI-500 sequencing platform (BGI-Shenzhen, Shenzhen, China). After quality control, 100bp pair-end clean reads corresponding to 30X sequencing coverage were acquired. Sequence alignment to the reference genome (hg19) and mutation calling were performed by applying the “best-practice” GATK workflow (21). Mutations were excluded based on the following criteria: 1) with a frequency higher than 0.01 in the ExAC (All and Asian), 1000Genome (All and Asian) and ESP6500 databases; 2) with minor allele frequency less than 10% or greater than 90%; 3) with a reference SNP ID (rs) number in dbSNP build 147. Structure variations in RC-K8 cell line were detected using Manta (22) and copy number variations were estimated by Control-FREEC (23).

### Transcriptome sequencing

Total RNA of RC-K8 was sequenced at BGI-Shenzhen using Illumina HiSeq 2000. Reads were aligned to the reference genome (Hg38) by STAR (two-pass approach) (24) for transcriptomic profiling (by Cufflinks) (25) and mutational detection (by GATK) (26), and also aligned to the reference transcriptome (Hg19) by SOAP2 and SOAPfusion (27) for detecting gene-fusion events. Gene expression levels were reported as fragments per kilobase of transcript per million mapped reads (FPKM).

### CRISPR-Cas9 screen with pooled lentiCRISPRv2 library

Following a previously described procedure (28), a cell line with stably expressed Cas9 (RC-K8-Cas9) was generated and 300 million RC-K8-Cas9 cells were subsequently transduced by the pooled lentiCRISPRv2 library (multiplicity of infection ∼0.3). One hundred-fifty million cells were collected 24 hours after the transduction as the baseline, and the remaining cells were cultured under 2.5µg/ml puromycin (Sigma, Darmstadt, Germany) selection for 10 days. At day-3, 7 and 14, respectively, 110, 60 and 60 million cells were collected, whereas 120 million cells were kept in culture after day-14 and treated with either JQ1 (400nM) (Tocris Bioscience, Abington, UK) or DMSO for two additional weeks. For each treatment, 60 million cells were collected at two time-points: day-21 and day-28. Genomic DNA was extracted from cells collected at different time points using the Blood & Tissue kit (Qiagen, Hilden, Germany). Guide sequences were PCR-amplified and sequenced at the Broad Institute using an Illumina NextSeq sequencer, following a reference protocol (28). The screening procedure was performed independently in two replicate experiments.

### Pre-processing of CRISPR-Cas9 screen data

A modified pre-processing step derived from the original protocol (count_spacers.py script) (28) was applied, trimming raw reads in FastQ format by the spacer sequences from both ends (3-prime: GTTTT and 5-prime: CGAAACACC) using CutAdapt (v1.8.3 Martin, 2011). One mismatch was allowed in the 5-prime sequence (parameter -e 0.12). The trimmed reads with a length of 18-21bp were mapped to the FASTA library of guide sequences (GeCKO v1/v2, Addgene.org) using Bowtie2 (29). To increase the specificity of read-count data, we mapped the library against to the human genome sequence and excluded guide sequences targeting non-protein coding regions (named with “mir-” or “let-” tags) or having alternative alignments (filtered by “XS” and “NM” tags in the bowtie2 result). After filtering, read-count data were normalized by the median count of each individual experiment.

### Gene essentiality estimation

The MAGeCK maximum-likelihood estimation (MLE) algorithm was applied to estimate the relative screening effects (beta scores or modified log fold-changes) in both negative and positive directions, i.e., depletion and enrichment. In the design-matrix of the MAGeCK analysis, samples were binary coded in different groups by the properties of time points, batches and drug treatments (Table S1). Samples from day-1 were labeled as the baseline (zero) in all grouping conditions. As a result, one beta score was calculated by MLE for each gene in each condition. In order to resolve the CRISPR screening effect with the time-course, beta-scores were fit by linear interpolation curves along the time-axis (k days). For every gene, given that day-1 is the baseline: *β*(1) = 0, the integral of beta-scores at a time point (k) was used as the proxy of the overall CRISPR screening effect: 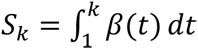. To normalise the integral scores, null distributions of scores were estimated based on a null set of read-count data generated by permutating the identities of single-guide RNA (sgRNA) sequences. Null scores were calculated by MAGeCK-MLE following the same procedure, and then fitted by Gaussian distributions at each time-point *k*. Subsequently, the original integral scores were standardized by the mean and standard deviation of the null distribution: 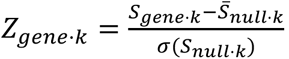.

To test the validity of the scoring method, two reference datasets (HT-29 (30) and CERES (31)) were used for benchmarking, i.e., the HT-29 dataset (colon cancer cell line, with multiple time points) and the CERES dataset (341 cancer cell lines, with one time point). The Pearson’s correlation coefficients between the estimated CRISPR effects and the general essentialities of the average CERES scores served as the figure-of-merit for overall performance at each time point. In addition, two previously published CRISPR screens were used to characterize essential DLBCL genes (10, 11). To be able to compare the results from different dataset, for each cell line, we also ranked the CRISPR scores for 16821 genes that were studied in all samples (10,11,31).

Gene set enrichment analysis (GSEA) was performed by applying the pre-ranked GSEA approach (32). The 50 hallmark gene sets (33) were used in GSEA, while weights were set to zero (classic method). 1000 times permutations were applied, and normalized enrichment scores (NES) were used for comparisons between datasets.

### Single-gene knockout experiments

Two to three sgRNA for each of the targeted genes and non-targeted controls were chosen from the lentiCRISPRv2 library and individually cloned into the plasmid backbone of the sgRNA library. Lentiviruses were subsequently produced in 293T cells for each selected sgRNA. In each single-gene knockout experiment, 3 million RC-K8-Cas9 cells were transduced and then selected by puromycin (5µg/ml). The growth of cells was monitored by counting the numbers of living cells at five time points (day-1, 3, 5, 9 and 14). DNA was extracted from cells collected at day-5 to verify the desired targeting (introduction of loss-of function insertions or deletions) of each selected sgRNA by SURVEYOR assay (Integrated DNA Technologies, Coralville, USA) (28) and sequencing.

### Cell viability assay

2×10^4^ cells were seeded in a volume of 100μL per well with vehicle or the indicated concentrations of drugs (nutlin-3, Santa Cruz Biotechnology, USA; chidamide, Chipscreen Biosciences, Shenzhen, China; metformin or SGC-30, Sigma-Aldrich). Metformin was dissolved in water, whereas the other drugs were dissolved in DMSO. 72h post-treatment, 20μl per well of CellTiter 96 AQueous One Solution Reagent (Promega, Madison, USA) was added. After incubation of the plates for 4h at 37°C, cell viability was measured by comparing the absorbance (A) at 450nm: A_treatment_/A_control_ ×100%. Each experiment was independently repeated at least three times.

### Statistics

P-values were calculated by Student’s t-test/ANOVA for quantitative comparisons, and χ^2^ or Fisher’s exact test for categorical comparisons.

### Data Sharing Statement

The CRISPR scores and read-counts for gRNA for the CRISPR screen are shown in Table S2 and Table S3, respectively. For raw data of library sequencing, WGS and RNAseq, please contact.

## Results

### Characterization of genome and transcriptome of RC-K8 cells

To characterize the genetic alterations in the RC-K8 cell line, we performed whole genome and transcriptome sequencing (WGS and RNA-seq) on DNA and RNA samples derived from this cell line. Based on the distribution of genomic sequence coverage, we discovered trisomy of chromosome 7 and gain of part of chromosomes 5, 13 and 20 (Figure S1) and estimated copy number gains of *MYC* and *NOTCH1* as well as amplification of *REL* (Table S2). In addition, RC-K8 cells were estimated to have copy number losses of *CD70*, *UBE2A* and two cohesion related genes *STAG2* and *SMC1A* (Table S2). We further identified 384 genes with non-synonymous mutations, including genes involved in DNA damage response and repair (*RAD21*, *TP63*, *TP73* and *XRCC6*), BCR/ NF-κB signaling (*TNFAIP3* and *NFKBIA*) and transcription factors important for B-cell development (*IKZF1* and *IKZF3*) (Table S2). Sequencing data showed that RC-K8 cell line harbors a largely normal *TP53* gene with a benign P72R polymorphism, but with a relatively high expression level of its negative regulator *MDM2* (FPKM: 46.55) as estimated by RNA-seq. Through WGS, we also identified structural variants including translocations involving *IGH* and *BCL6* (with different partners respectively) as well as a balanced translocation between chromosomes 22 and 6 in RC-K8 (Figure S2A), which resulted in a C-terminal truncation of *EP300* that has been previously reported as EP300ΔC1047 (20, 34). RNA-seq analysis further demonstrated dominant allelic expression of the truncated form of *EP300* (Figure S2B), which is encoded by exons 1-17 of *EP300* and fused to 25 AAs encoded by intronic sequences of *BCKDHB* gene on chromosome 6, resulting in loss of two critical domains of this protein, i.e., the bromodomain and HAT-domain (20).

### Essential genes and pathways in RC-K8 cells

To investigate the cancer dependencies related to the genetic background of RC-K8 cell line, we performed a functional genetic screen by introducing the genome-wide pooled CRISPR-Cas9 knock-out library (detailed in Methods). A Cas9-expressing RC-K8 cell line (RC-K8-Cas9) was first generated and subsequently transduced by the pooled lentiCRISPRv2 library, which targets 19,050 human genes with more than 100,000 unique sgRNAs. By deep-sequencing the amplified sgRNA library, we identified depleted gRNAs caused by dropouts of cells bearing the related genetic perturbations, which reflects the importance of the targeted genes for cell growth or proliferation.

We next tested our optimized MAGeCK-based scoring approach (detailed in Methods) on the HT-29 dataset (a colon cancer cell line, with multiple time points, day 3 to day 25) (30) and the CERES dataset (341 cancer cell lines, with one time point) (31). The correlations between the standardized CRISPR scores, i.e., standardized accumulative beta-scores for HT-29 cells at different time points and the average of CERES scores, representing the general essentialities, showed an increasing trend along the time course – correlation coefficients ranged from 0.56 (at day-7) to 0.78 (at day-25). The correlations are generally stronger than using the reported CRISPR scores calculated from pairwise comparisons between the baseline and individual time-points, suggesting an improved estimation of gene essentialities by our scoring method.

We then applied the novel scoring system to estimate the relative effects of gene knockouts during the 28 days’ time-course of the CRISPR screen on RC-K8 cells (Figure 1A). Based on both the read-count distribution and the standardized CRISPR scores, we observed no significant depletion at day-3 after transduction (Figure 1B). In contrast, at day-7 we started to observe a rapid depletion of cells bearing knockouts of core-fitness genes related to ribosome, spliceosome, proteasome and cell-cycle regulation (7,8,18,32,35–37) (Figure 1C). At day-7, the top candidates for core-fitness genes also included cell-type specific essentialities (Table S2). Several pan-DLBCL essential genes, as estimated from the previous two screens (10, 11), were detected early at day-7, including *FOXO1*, *IRF4* and *SF3B1*, while some appeared later than day-14, for example *TAF1*, *MLL2*, *RHOA* and *YY1*. More specific for the RC-K8 cells, which carry a largely normal *TP53* gene locus, *MDM2* was one of the most significant essential genes detected at day-7, demonstrating its critical role in suppressing TP53-induced cell death. *CCND3*, which is associated with both TP53-signaling and cell-cycle regulation pathways, exhibited essentiality and high specificity to the cell line. Similarly, *CREBBP* was also identified at day-7 as one of the top-hits, showing its essentiality in the context *EP300* loss-of-function.

**Figure 1.**
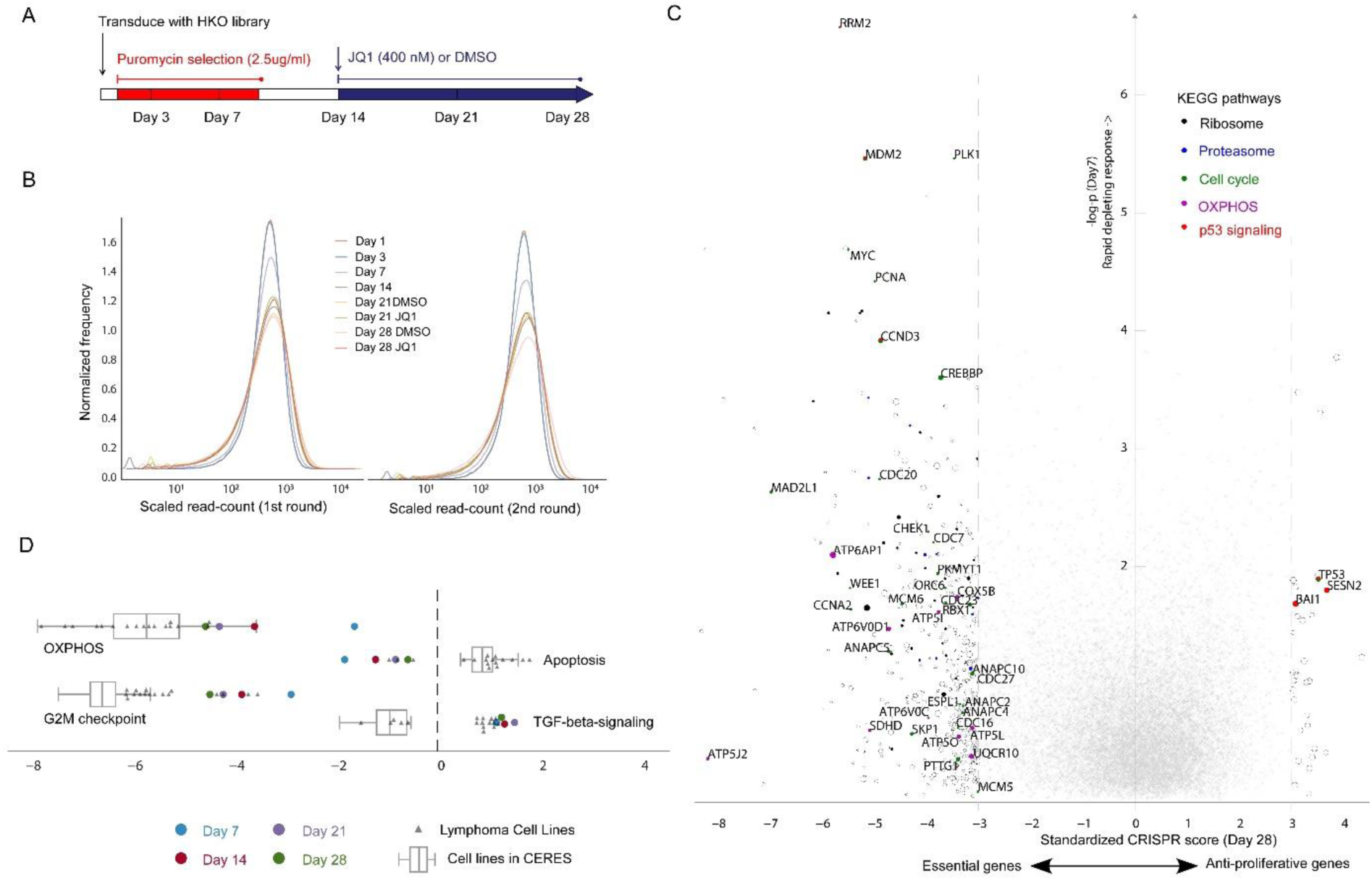
Design and result of the CRISPR-Cas9 loss-of-function screen in RC-K8. **A)** Time points and drug treatments. The lentiCRISPR GeCKOv2 library was transduced at day-zero. Puromycin selection of successful transduction was applied between day-1 and day-10. Drug treatments (JQ1 and DMSO vehicle) were applied after the mid-point day-14. Cells were harvested at the six time-points for sequencing, including day-1 as the baseline. **B)** Distribution of read-counts of guide sequences (rescaled and centered by the median of 1000). Compared to the baseline and the later time points, the distribution at day-7 is distinctive, which has been used for detecting essential genes that caused rapid depletion. **C)** Dropout effects of gene knockouts measured at different time points; y-axis: significance of rapid responses to genetic perturbations observed at day-7; x-axis: standardized (z-transformed) CRISPR-score reflecting overall effects at day-28. Circles: genes with overall scores <-3.0 (essential genes) or >3.0 (anti-proliferative genes); Circle sizes are scaled by the average CERES score reflecting general essentiality in cancer cells, where smaller circles indicate cell-specific dependencies, such as *MDM2*, *CREBBP* and *CCND3*. Genes involved in the five indicated KEGG pathways are filled with colors. **D)** Gene-set enrichment analysis (GSEA) of four selected hallmark pathways. Normalized enrichment scores (NES) based on the screening results of RC-K8 cells at four time points (colored dots) were compared to NES based on previously reported screens from B-cell lymphoma cell lines (grey triangles) and 341 cancer cell lines from the CERES data (box-plot). OXPHOS pathway was the most differentially enriched between the early and late time points in RC-K8. Compared to other cell lines, RC-K8 cells showed stronger dependency on the apoptosis pathway, in which the major contributor is *CREBBP* that exhibited cell-specific essentiality. By comparing enrichment scores between lymphoma cell lines and other cancer cell lines in CERES, several pathways were significantly differential, e.g., the G2M-checkpoint (t-statistic >13.07) and TGF-beta signaling (effects in different directions). The comparisons of all 50 hallmark pathways are shown in Figure S4.

On the axis of positive selection, the effects of knocking out growth-suppressing genes were generally accumulative. We observed a significant anti-proliferative effect of the classical tumor suppressor *TP53* and several associated regulators/effectors, such as *SESN2* and *BAI1* (Figure 1C). Knockouts of some “cancer drivers” of DLBCL, including *BCL2*, *BCL6*, *NFKBIA* and *CD70*, showed only modest promoting effects of cell proliferation (Table S2).

The standardized CRISPR scores at day-7 exhibited a weak correlation (Pearson’s r < 0.25) with the average CERES scores. On the contrary, the cumulative effects at later time points correlated stronger with the CERES scores, with highest correlation at day-28 (Pearson’s r > 0.48) (Figure S3, Table S2). We surmise that rapid depletions were caused by genetic lethality, whereas delayed dropouts were associated with impaired cellular function or fitness. We hypothesize that essential genes exhibiting different rates of depletion may participate in complementary pathways. To test this hypothesis, based on the ranks of CRISPR scores, we applied gene set enrichment analysis against 50 hallmark pathways (Figure S4, Table S4). We observed that the most differentially enriched pathways between early and late time points (day-7 vs. day-28), were apoptosis and oxidative phosphorylation (OXPHOS) pathways (Figure 1D). Compared to normalized enrichment scores of the 341 cell lines from the CERES data set, the rapid depletion of apoptosis related genes showed cell-type specificity, where *BCL2L1*, *CREBBP* and *WEE1* were among the most significant genes. On the other hand, depletions associated with OXPHOS genes showed gradual increase during the time-course, suggesting that growth suppression caused by the interrupted energy metabolism is slow and cumulative

The data also allowed us to investigate, within the same analysis, the relative effects associated with JQ1 (a BET inhibitor) treatment during the screening process (Table S2). BET inhibition by JQ1 has been suggested to trigger cell-cycle arrest followed by apoptosis or senescence (35), and we indeed observed a long-term growth-suppressive effect in the JQ1-treated cells. We found no significant enrichment of positively selected genes (those possibly causing resistance to the JQ1 treatment). Nonetheless, several negatively selected genes were found to be associated with the MAPK signaling pathway, including *PAK2*, *LAMTOR3* and *MAPKAPK2*. Notably, the top candidate gene, *PAK2*, was non-essential in RC-K8 cells in general, but was selectively depleted in the JQ1-treated cells. This may indicate that the loss of *PAK2* could sensitize RC-K8 cells to JQ1 treatment.

We validated several top-ranking genes (essential genes *MDM2* and *CREBBP*, tumor suppressor gene *TP53*) from the genome-wide screening by single-knockout experiments (Figure S5A-D). We further performed cell viability assays by targeted inhibition of top candidate genes using small molecular drugs. We confirmed the rapid and complete responsiveness to *MDM2* inhibitions by nutlin-3 (Figure S6A). Additionally, when given a high concentration of metformin (a widely-used drug that inhibits OXPHOS), the tested lymphoma cells, including RC-K8, responded to the treatment (Figure S6B), suggesting a dependency on the OXPHOS pathway.

### DLBCL cells with either EP300 or CREBBP mutations are sensitive to HAT domain inhibitor

By re-analyzing a Chinese DLBCL cohort (5), we observed that mutations were more frequently identified in *CREBBP* than that in *EP300* (11.7% vs. 3.3% of 275 samples, Figure 2A, Table S5), which is in agreement with previous studies (12, 36). Notably, over 70% of these mutations affected the HAT-domain (Figure 2A). In most of the cases, we observed either *CREBBP* or *EP300* mutations, however in several DLBCL samples, we identified mutations in both genes, although the oncogenic role of these mutations remains unknown in some cases (Table S5).

**Figure 2.**
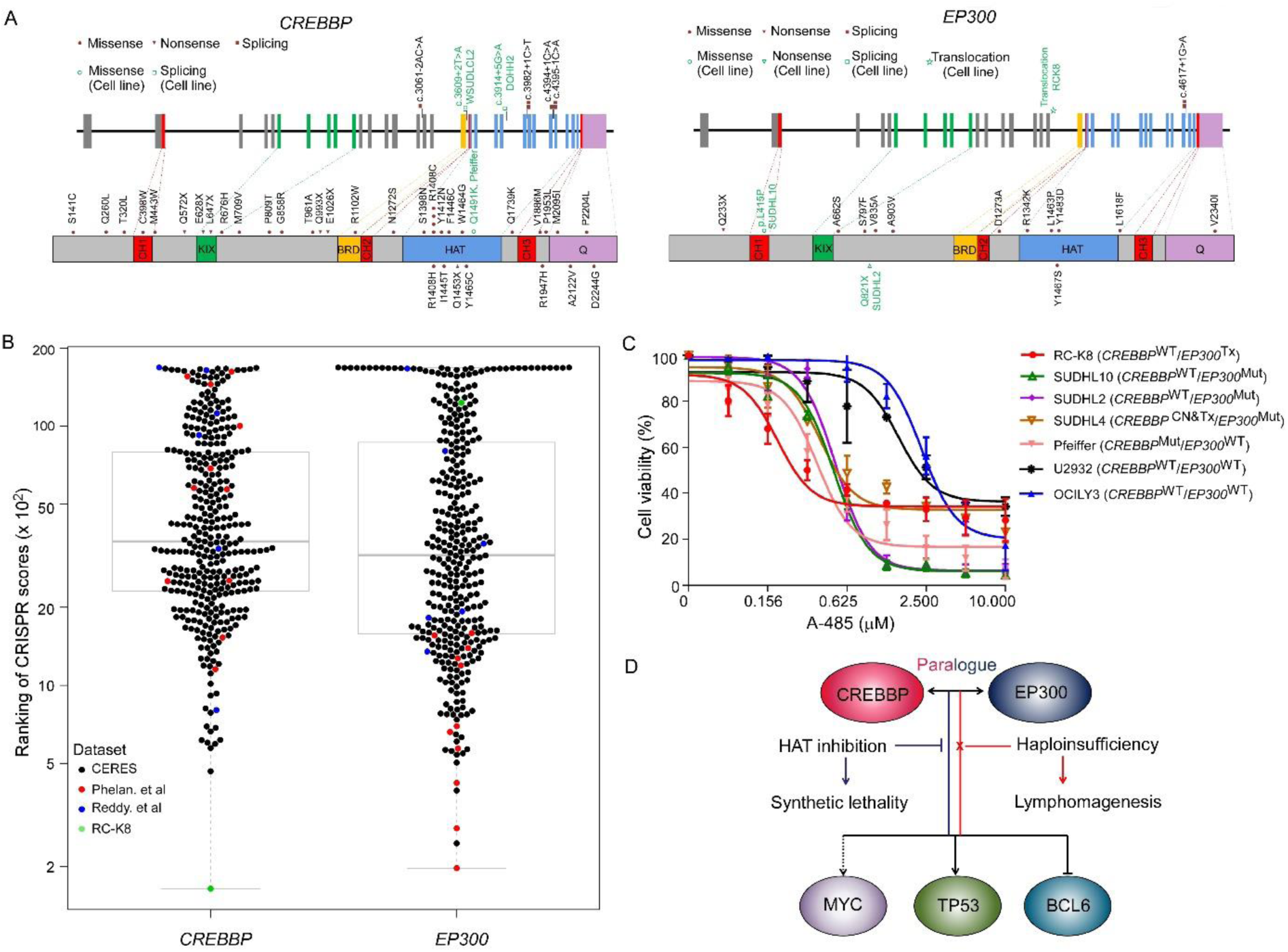
DLBCL cells with genetic alterations in *EP300* and *CREBBP* are sensitive to a HAT domain inhibitor. **A)** Somatic mutations in *CREBBP* and *EP300* identified in the Chinese DLBCL cohort (5), the RC-K8 cells and other cell lines included by previous CRISPR screen or tested by A-485 in this study. **B)** Ranking of CRISPR scores of CREBBP and EP300 from different datasets. The middle boxes represent the middle 50% of values for each group with a mid-line as the median value. The bars outside the boxes represent the 1.5 interquartile ranges outside of the boxes. **C)** Cell viability tests of targeted inhibitions of HAT-domain by small molecule A-485. Error-bars: standard deviation (n=3 replicates). Compared to cell lines (OCILY3 and U2932) with wild-type HATs, the five HAT-mutant cell lines displayed higher sensitivity to the inhibition, with RC-K8 cells exhibiting the lowest IC50 value. CN: copy number loss; Tx: translocation. **D)** Potential mechanism of HAT-inhibition in HAT-deficient cells: haploinsufficiency of *CREBBP* or *EP300* promotes lymphomagenesis via dysregulation of BCL6 and TP53 (18); the deficiency develops dependency on the remaining HAT function at the same time. Targeted inhibition of HAT could causes synthetic lethality through abrogation of MYC (19).

Based on re-evaluating and ranking the CRISPR scores of the reference datasets, *CREBBP* and *EP300* seemed to be essential in a small set of cell lines (Figure 2B). Among the DLBCL cell lines analyzed, *CREBBP* is almost uniquely essential in the RC-K8 cells, whereas *EP300* is more frequently identified as an essential gene (Figure 2B and Table S6), which may reflect the different mutational rates in these genes in DLBCL patients. Importantly, three GCB-like DLBCL cell lines that were highly sensitive for *EP300* knockout harbored mutations (DOHH2 and WSUDLCL2) or CNV loss (SUDHL5) in *CREBBP* (Figure 2B and Table S6). Notably, one GCB cell line (SUDHL4) had genetic alterations in both *CREBBP* and *EP300* genes, and it was more sensitive to *EP300* knockout (Table S6). This may be due to the difference in the nature of genetic changes in this cell line: a translocation and a CNV loss were identified in *CREBBP*, but only a missense mutation was observed in *EP300*.

We next performed cell viability tests using a newly reported catalytic inhibitor (A-485) that targets the HAT-domain of both *CREBBP* and *EP300* (37). RC-K8 cells responded to the A-485 treatment, with the lowest IC50 value (0.42 µM) compared to six other DLBCL cell lines tested (Figure 2C and Table S6). In general, the five cell lines with *CREBBP*- and/or *EP300*-mutations showed a higher sensitivity than the two cell lines wildtype for these genes (OCILY3 and U2932) (Figure S7), suggesting that dependency of the remaining HAT function is a druggable vulnerability in DLBCL cells with either *CREBBP* or *EP300* genetic alterations (Figure 2D).

It has previously been reported that the bromodomains are also critical to sustain the proliferation of lymphoma cells via epigenetic regulation. However, the efficacy of the pan-BET family inhibitor JQ1 is unlikely to be related to the HATs due to its low affinity to the bromodomain of *CREBBP*/*EP300* (38). Therefore, we tested the *CREBBP*/*EP300*-specific bromodomain inhibitor CBP30 and the RC-K8 cells showed the lowest sensitivity compared to four other cell lines (Figure S6C).

Deficiencies in HATs may change the balance of histone acetylation and alter chromatin structure. Therefore, counteracting the epigenetic changes by inhibiting the activity of histone deacetylases (HDACs) creates a therapeutic option. Reportedly, RC-K8 cells show a low level of histone acetylation due to the deficiency of *EP300* (36). However, we observed no significant essentiality of HDACs in the CRISPR screen, except *HDAC3*, that is deemed a core-essential gene in most cancer cells (Table S2). Furthermore, in our cell viability tests, the pan-HDAC inhibitor (Chidamide) showed the least sensitivity in RC-K8 cell line compared to four other DLBCL cell lines (Figure S6D). These results may suggest that targeting HATs, instead of HDACs, could be more effective in the treatment of B-cell lymphomas, especially in those with HAT-deficiency due to genetic alterations in *CREBBP* and *EP300*.

## Discussions

In this study, we performed a genome-wide CRISPR-Cas9 loss-of-function screening and identified *CREBBP* as a vulnerability specific to a DLBCL cell line RC-K8, which harbored a translocation that disrupts the *EP300* gene. *CREBBP* is one of the most frequently mutated genes in non-Hodgkin lymphoma (NHL) and usually plays a tumor-suppressing role through epigenetic regulation (18). Such dependency on *CREBBP* can be best explained by a synthetic lethality associated with the loss-of-function of its paralogue *EP300* (19) (Figure 2D). As reported in multiple myeloma (39), the bromodomain of *CREBBP* is essential for regulating the pan-DLBCL essential gene *IRF4*. Although *IRF4* also exhibited essentiality in our CRISPR screen with the RC-K8 cells, the bromodomain inhibitor CBP30 did not show efficacy. On the contrary, A-485, a potent catalytic inhibitor of HAT-domains (37), showed high efficacy in this *EP300*-deficient cell line. The results suggested that the HAT-domain of *CREBBP* is linked to the essentiality in RC-K8 cells. By re-evaluating the previously published CRISPR-Cas9 screens from a pan-cancer study and sets of DLBCL cell lines, we also identified a genetic essentiality of *EP300* or *CREBBP* gene in a small subset of cancer cell lines, including a few DLBCL cell lines that are dependent on *EP300* and carry *CREBBP* mutation, copy number loss or translocation. We further observed differences in IC50 values of A-485 in seven lymphoma cell lines, indicating a potential correlation between the deficiency of *CREBBP/EP300* and the sensitivity to the HAT-inhibitor. Our results thus suggest that targeting the remaining HAT function may hold therapeutic potential for B-cell lymphomas with deficiency in either *CREBBP* or *EP300*.

The genetic features of the RC-K8 cell line showed some similarity to the BN2 (based on *BCL6* fusions and *NOTCH1* CNV changes) (4) or the C1 (based on *BCL6* structural variants and mutations of *NOTCH* signaling pathway components) molecular subtype (3), but also have distinct genetic features. We furthermore noted that it resembled what we have discovered previously in DLBCL samples associated with HBV infection (5), with translocations in *BCL6*, copy number changes in *NOTCH1* and *CD70*, as well as mutations in *ZFP36L1*, *SGK1*, *IKZF3*, *TP63* and *TP73*. It has been shown that the HBV protein HBx can directly interact with *CREBBP*/*EP300* and facilitate the recruitment of the complex onto CREB-responsive promoters, upregulating downstream oncogenes (40). This was supported by the observation that the *CREBBP*-targeted genes were significantly upregulated in HBsAg^+^ tumors compared to HBsAg^-^ tumors (Figure S8). Targeting the HAT-functions of *CREBBP/EP300* can thus be a new direction in developing effective treatment for HBV-associated DLBCL patients, who usually have a poor response to the current therapy (5).

In addition to *CREBBP*, we also identified a cell line-specific essentiality of *MDM2* in RC-K8, which has an increased level of expression of *MDM2* and a largely unaffected *TP53* gene. Accordingly, the screen also showed a strong positive selection of *TP53* in RC-K8 cells. *TP53* is mutated in about 50% of human cancers and is one of most studied tumor suppressors (41). In other tumors, including the majority of the DLBCLs however, *TP53* is in its wild-type form, and targeting the negative regulators for *TP53*, such as *MDM2*, may be a promising approach (42). When re-analyzing the CERES dataset, we indeed observed that cancer cell lines with wild-type *TP53* in general are more dependent on *MDM2* and *MDM4* (Figure S9). A recent CRISPR-Cas9 screen has also identified druggable dependencies in *TP53* wild-type Ewing sarcoma, including *MDM2*, *MDM4*, *USP7* and *PPM1D* (43).

RC-K8 cells have a normal functional TP53, which is capable of triggering DNA-damage responses to the CRISPR-Cas9 editing during the screening process (44). Consequently, the calling of essential genes through a depleting effect might be affected by TP53-induced apoptosis or cell-cycle arrest. We therefore used a time course experimental design to distinguish between essentiality and DNA-damage response, as cells carrying dispensable knockouts may recover from the perturbation and then grow back. We observed that many such genes exhibited a rapid depleting effect but were not ultimately essential (Table S2). During the time-course of the CRISPR screen, we also observed a delayed depleting effect from many essential genes and pathways, notably the OXPHOS-related genes. Such delayed essentialities of metabolic pathways are repeatedly observed in CRISPR-based genetic screens (30, 45). One possible reason for the delayed depletion is the poor correlation between mRNA and protein expression, which has been observed in proteomics studies (46). Thus, highly expressed genes may provide a buffer to the CRISPR-induced perturbations and cause false negative results in the essentiality calling. The time-course information, analyzed by our optimized method, may help to detect such delayed essentialities in cancer cells.

By introducing drug treatments to the CRISPR screen, we identified *PAK2* as a potentially essential gene for survival of RC-K8 cells under the stress induced by the pan-BET inhibitor JQ1. The result suggested that dual blockage of BET-family proteins and MAPK-signaling pathway could be an effective treatment of not only colorectal (47) or ovarian cancer (48), but also B-cell lymphomas. Further functional validation will, however, be required to confirm the screen result.

In conclusion, our unbiased, genome-wide, time course-based CRISPR-Cas9 screen revealed a number of cell line-specific vulnerability, such as *CREBBP* and *MDM2*, and a delayed metabolic dependency of OXPHOS genes. By re-analyzing the previously published CRISPR-Cas9 screens, we also identified a genetic essentiality of *EP300* in additional DLBCL cell lines. Considering the high mutation rates of *CREBBP* and *EP300* in DLBCL and follicular lymphoma (FL), two most common types of NHL, and the prognostic value of these two genes in FL (M7-FLIPI) (49), the findings of our study provide insights for development of more effective targeted therapies as well as novel combination treatments that may benefit a large group of patients. It is furthermore important to point out that DLBCL is a highly heterozygous disease and each tumor carries unique combination of genetic alterations, affecting multiple functional pathways. Characterization of “core fitness” gene for cancer cells and “pan-DLBCL” essential genes might be informative for prioritization of therapeutic targets (9), for each cell line or individual/subgroup of patients however, integration of genetic/transcriptomic data with unbiased functional screen will still be needed to identify the most effective targeted therapy.

## Acknowledgements

This work was supported by the Swedish Cancer Society, the Swedish Research Council, the European Research Council (RNAEDIT-649019), the Swedish Childhood Cancer Fund, the Chinese Natural Science Foundation (81670184), STINT (Joint China-Sweden mobility program), Radiumhemmets and the Center for Innovative Medicine (CIMED). B. Zhang was supported by a Cancerfonden fellowship. The computational resource was provided by SNIC through Uppsala Multidisciplinary Center for Advanced Computational Science (UPPMAX) under project SNIC 2018/8-70. The authors appreciate Prof. Zubarev’s lab at KI for sharing anti-cancer chemicals and Dr. X. Lu from Chipscreen for providing chidamide.

## Author contributions

B. Z., X. Y., and M. N. analyzed and interpreted the screen data; L. D., M. N. and X. S. performed the experiments; W. R. analyzed the mutations in the DLBCL cohort; J. J. helped with the screening experiment design and analysis; X. Y., D. L, W. R. and K. W. analyzed the WGS data; F. Z. and Q. P. -H. supervised the study; B. Z., M. N., and Q.P.-H. wrote the manuscript.

## References

1. Alizadeh AA, Eisen MB, Davis RE, Ma C, Lossos IS, Rosenwald A, et al. Distinct types of diffuse large B-cell lymphoma identified by gene expression profiling. Nature 2000;403:503–11

2. Basso K, Dalla-Favera R. Germinal centres and B cell lymphomagenesis. Nat Rev Immunol 2015;15:172–84

3. Chapuy B, Stewart C, Dunford AJ, Kim J, Kamburov A, Redd RA, et al. Molecular subtypes of diffuse large B cell lymphoma are associated with distinct pathogenic mechanisms and outcomes. Nat Med 2018;24:679–90

4. Schmitz R, Wright GW, Huang DW, Johnson CA, Phelan JD, Wang JQ, et al. Genetics and Pathogenesis of Diffuse Large B-Cell Lymphoma. N Engl J Med 2018;378:1396–407

5. Ren W, Ye X, Su H, Li W, Liu D, Pirmoradian M, et al. Genetic landscape of hepatitis B virus-associated diffuse large B-cell lymphoma. Blood 2018;131:2670–81

6. Shalem O, Sanjana NE, Zhang F. High-throughput functional genomics using CRISPR-Cas9. Nat Rev Genet 2015;16:299–311

7. Wang T, Birsoy K, Hughes NW, Krupczak KM, Post Y, Wei JJ, et al. Identification and characterization of essential genes in the human genome. Science 2015;350:1096–101

8. Hart T, Chandrashekhar M, Aregger M, Steinhart Z, Brown KR, MacLeod G, et al. High-Resolution CRISPR Screens Reveal Fitness Genes and Genotype-Specific Cancer Liabilities. Cell 2015;163:1515–26

9. Behan FM, Iorio F, Picco G, Goncalves E, Beaver CM, Migliardi G, et al. Prioritization of cancer therapeutic targets using CRISPR-Cas9 screens. Nature 2019;568:511–6

10. Reddy A, Zhang J, Davis NS, Moffitt AB, Love CL, Waldrop A, et al. Genetic and Functional Drivers of Diffuse Large B Cell Lymphoma. Cell 2017;171:481–94 e15

11. Phelan JD, Young RM, Webster DE, Roulland S, Wright GW, Kasbekar M, et al. A multiprotein supercomplex controlling oncogenic signalling in lymphoma. Nature 2018;560:387–91

12. Jiang Y, Ortega-Molina A, Geng H, Ying HY, Hatzi K, Parsa S, et al. CREBBP Inactivation Promotes the Development of HDAC3-Dependent Lymphomas. Cancer Discov 2017;7:38–53

13. Pasqualucci L, Dalla-Favera R. Genetics of diffuse large B-cell lymphoma. Blood 2018;131:2307–19

14. Rosenquist R, Bea S, Du MQ, Nadel B, Pan-Hammarstrom Q. Genetic landscape and deregulated pathways in B-cell lymphoid malignancies. J Intern Med 2017;282:371–94

15. Kung AL, Rebel VI, Bronson RT, Ch’ng LE, Sieff CA, Livingston DM, et al. Gene dose-dependent control of hematopoiesis and hematologic tumor suppression by CBP. Genes Dev 2000;14:272–7

16. Vo N, Goodman RH. CREB-binding protein and p300 in transcriptional regulation. J Biol Chem 2001;276:13505–8

17. Pasqualucci L, Dominguez-Sola D, Chiarenza A, Fabbri G, Grunn A, Trifonov V, et al. Inactivating mutations of acetyltransferase genes in B-cell lymphoma. Nature 2011;471:189–95

18. Zhang J, Vlasevska S, Wells VA, Nataraj S, Holmes AB, Duval R, et al. The CREBBP Acetyltransferase Is a Haploinsufficient Tumor Suppressor in B-cell Lymphoma. Cancer Discov 2017;7:322–37

19. Ogiwara H, Sasaki M, Mitachi T, Oike T, Higuchi S, Tominaga Y, et al. Targeting p300 Addiction in CBP-Deficient Cancers Causes Synthetic Lethality by Apoptotic Cell Death due to Abrogation of MYC Expression. Cancer Discov 2016;6:430–45

20. Garbati MR, Thompson RC, Haery L, Gilmore TD. A rearranged EP300 gene in the human B-cell lymphoma cell line RC-K8 encodes a disabled transcriptional co-activator that contributes to cell growth and oncogenicity. Cancer Lett 2011;302:76–83

21. Van der Auwera GA, Carneiro MO, Hartl C, Poplin R, Del Angel G, Levy-Moonshine A, et al. From FastQ data to high confidence variant calls: the Genome Analysis Toolkit best practices pipeline. Curr Protoc Bioinformatics 2013;43:11 0 1-33

22. Chen X, Schulz-Trieglaff O, Shaw R, Barnes B, Schlesinger F, Kallberg M, et al. Manta: rapid detection of structural variants and indels for germline and cancer sequencing applications. Bioinformatics 2016;32:1220–2

23. Boeva V, Popova T, Bleakley K, Chiche P, Cappo J, Schleiermacher G, et al. Control-FREEC: a tool for assessing copy number and allelic content using next-generation sequencing data. Bioinformatics 2012;28:423–5

24. Dobin A, Gingeras TR. Mapping RNA-seq Reads with STAR. Curr Protoc Bioinformatics 2015;51:11 4 1-9

25. Trapnell C, Williams BA, Pertea G, Mortazavi A, Kwan G, van Baren MJ, et al. Transcript assembly and quantification by RNA-Seq reveals unannotated transcripts and isoform switching during cell differentiation. Nat Biotechnol 2010;28:511–5

26. DePristo MA, Banks E, Poplin R, Garimella KV, Maguire JR, Hartl C, et al. A framework for variation discovery and genotyping using next-generation DNA sequencing data. Nat Genet 2011;43:491–8

27. Jia W, Qiu K, He M, Song P, Zhou Q, Zhou F, et al. SOAPfuse: an algorithm for identifying fusion transcripts from paired-end RNA-Seq data. Genome Biol 2013;14:R12

28. Joung J, Konermann S, Gootenberg JS, Abudayyeh OO, Platt RJ, Brigham MD, et al. Genome-scale CRISPR-Cas9 knockout and transcriptional activation screening. Nat Protoc 2017;12:828–63

29. Langmead B, Salzberg SL. Fast gapped-read alignment with Bowtie 2. Nat Methods 2012;9:357–9

30. Tzelepis K, Koike-Yusa H, De Braekeleer E, Li Y, Metzakopian E, Dovey OM, et al. A CRISPR Dropout Screen Identifies Genetic Vulnerabilities and Therapeutic Targets in Acute Myeloid Leukemia. Cell Rep 2016;17:1193–205

31. Meyers RM, Bryan JG, McFarland JM, Weir BA, Sizemore AE, Xu H, et al. Computational correction of copy number effect improves specificity of CRISPR-Cas9 essentiality screens in cancer cells. Nat Genet 2017;49:1779–84

32. Subramanian A, Tamayo P, Mootha VK, Mukherjee S, Ebert BL, Gillette MA, et al. Gene set enrichment analysis: a knowledge-based approach for interpreting genome-wide expression profiles. Proc Natl Acad Sci U S A 2005;102:15545–50

33. Liberzon A, Birger C, Thorvaldsdottir H, Ghandi M, Mesirov JP, Tamayo P. The Molecular Signatures Database (MSigDB) hallmark gene set collection. Cell Syst 2015;1:417–25

34. Garbati MR, Alco G, Gilmore TD. Histone acetyltransferase p300 is a coactivator for transcription factor REL and is C-terminally truncated in the human diffuse large B-cell lymphoma cell line RC-K8. Cancer Lett 2010;291:237–45

35. Trabucco SE, Gerstein RM, Evens AM, Bradner JE, Shultz LD, Greiner DL, et al. Inhibition of bromodomain proteins for the treatment of human diffuse large B-cell lymphoma. Clin Cancer Res 2015;21:113–22

36. Hashwah H, Schmid CA, Kasser S, Bertram K, Stelling A, Manz MG, et al. Inactivation of CREBBP expands the germinal center B cell compartment, down-regulates MHCII expression and promotes DLBCL growth. Proc Natl Acad Sci U S A 2017;114:9701–6

37. Lasko LM, Jakob CG, Edalji RP, Qiu W, Montgomery D, Digiammarino EL, et al. Discovery of a selective catalytic p300/CBP inhibitor that targets lineage-specific tumours. Nature 2017;550:128–32

38. Filippakopoulos P, Qi J, Picaud S, Shen Y, Smith WB, Fedorov O, et al. Selective inhibition of BET bromodomains. Nature 2010;468:1067–73

39. Conery AR, Centore RC, Neiss A, Keller PJ, Joshi S, Spillane KL, et al. Bromodomain inhibition of the transcriptional coactivators CBP/EP300 as a therapeutic strategy to target the IRF4 network in multiple myeloma. Elife 2016;5

40. Neuveut C, Wei Y, Buendia MA. Mechanisms of HBV-related hepatocarcinogenesis. J Hepatol 2010;52:594–604

41. Kandoth C, McLellan MD, Vandin F, Ye K, Niu B, Lu C, et al. Mutational landscape and significance across 12 major cancer types. Nature 2013;502:333–9

42. Wade M, Li YC, Wahl GM. MDM2, MDMX and p53 in oncogenesis and cancer therapy. Nat Rev Cancer 2013;13:83–96

43. Stolte B, Iniguez AB, Dharia NV, Robichaud AL, Conway AS, Morgan AM, et al. Genome-scale CRISPR-Cas9 screen identifies druggable dependencies in TP53 wild-type Ewing sarcoma. J Exp Med 2018;215:2137–55

44. Haapaniemi E, Botla S, Persson J, Schmierer B, Taipale J. CRISPR-Cas9 genome editing induces a p53-mediated DNA damage response. Nat Med 2018;24:927–30

45. Li W, Xu H, Xiao T, Cong L, Love MI, Zhang F, et al. MAGeCK enables robust identification of essential genes from genome-scale CRISPR/Cas9 knockout screens. Genome Biol 2014;15:554

46. Mertins P, Mani DR, Ruggles KV, Gillette MA, Clauser KR, Wang P, et al. Proteogenomics connects somatic mutations to signalling in breast cancer. Nature 2016;534:55–62

47. Togel L, Nightingale R, Chueh AC, Jayachandran A, Tran H, Phesse T, et al. Dual Targeting of Bromodomain and Extraterminal Domain Proteins, and WNT or MAPK Signaling, Inhibits c-MYC Expression and Proliferation of Colorectal Cancer Cells. Mol Cancer Ther 2016;15:1217–26

48. Jing Y, Zhang Z, Ma P, An S, Shen Y, Zhu L, et al. Concomitant BET and MAPK blockade for effective treatment of ovarian cancer. Oncotarget 2016;7:2545–54

49. Pastore A, Jurinovic V, Kridel R, Hoster E, Staiger AM, Szczepanowski M, et al. Integration of gene mutations in risk prognostication for patients receiving first-line immunochemotherapy for follicular lymphoma: a retrospective analysis of a prospective clinical trial and validation in a population-based registry. The Lancet Oncology 2015;16:1111–22

