## supplemental figures for "Essentiality of *CREBBP* in *EP300* truncated B-cell lymphoma revealed by genome-wide CRISPR-Cas9 screen"

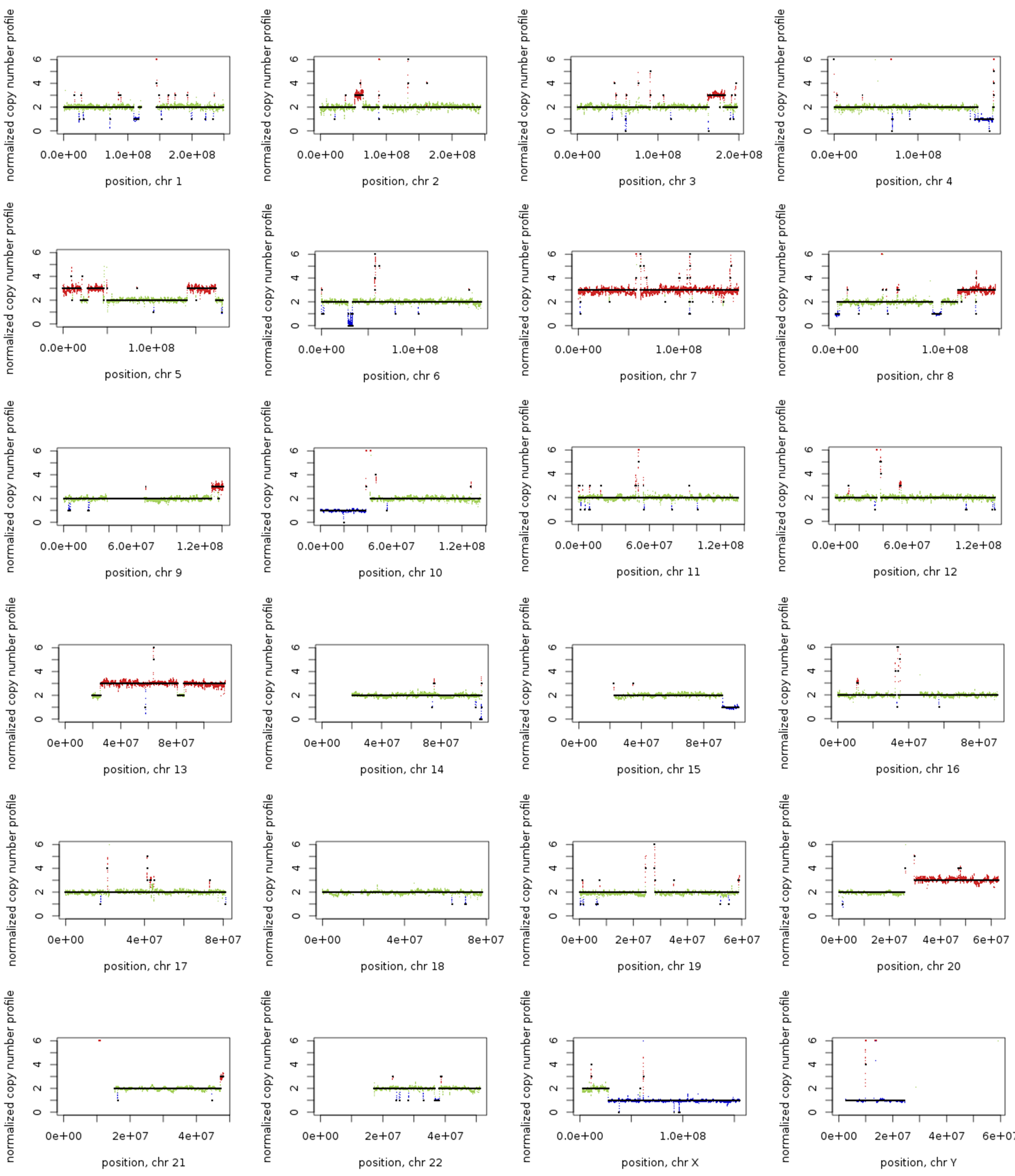

**Figure S1 | Overview of copy number variations (CNVs) in RC-K8 cell line.** CNVs of genomic regions were analysed by using the WGS data and the Control-FREEC tool (1). Whole chromosome or large part of chromosome gains of chromosomes 7 , 5, 13 and 20 were observed. Detailed information for CNVs at gene level is shown in Table S2.

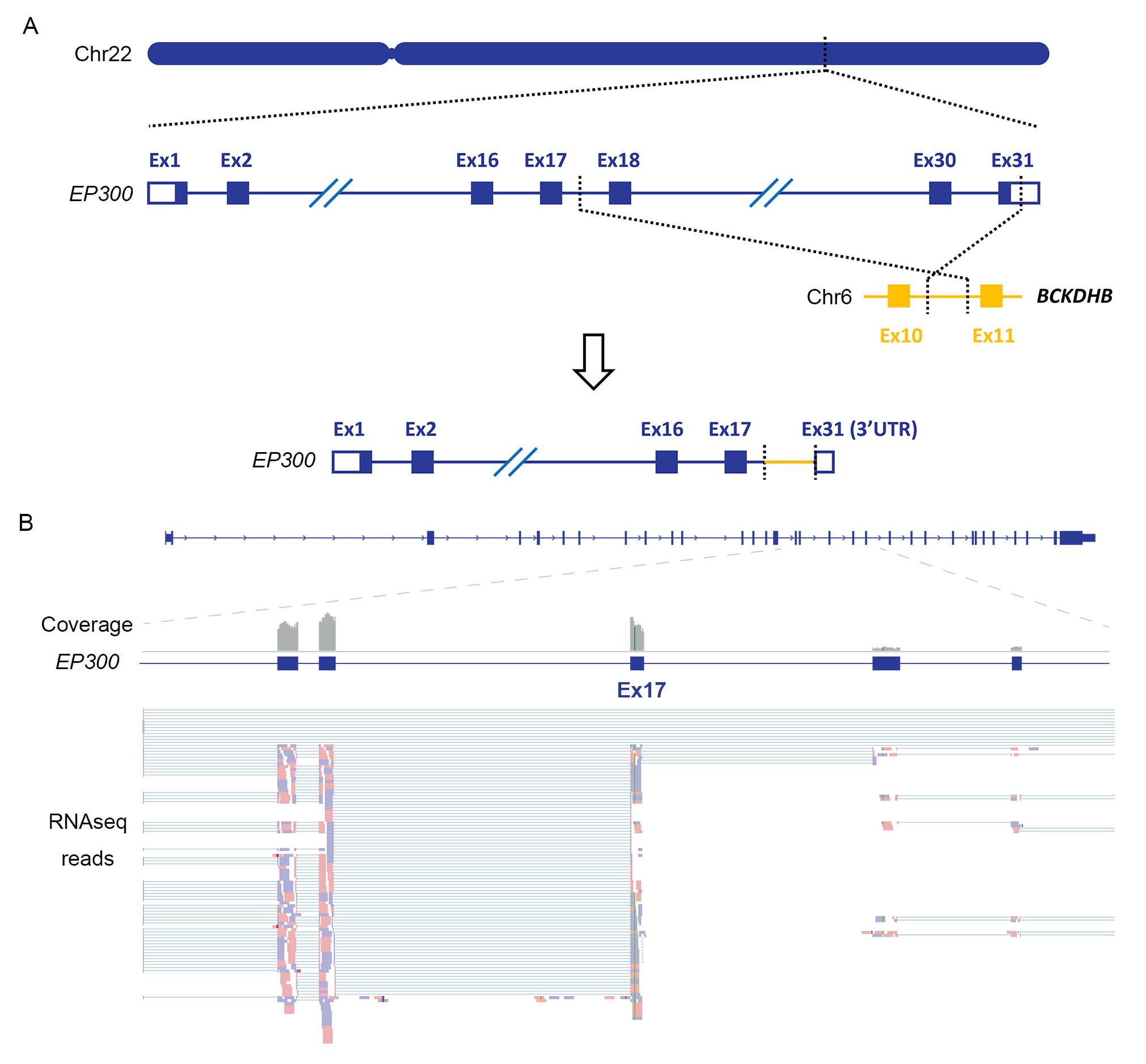

**Figure S2 | Identification of the *EP300* translocation in RC-K8 cell line by WGS and RNAseq.** A) Balanced translocation identified by WGS, involving *EP300* on chromosomal 22 and *BCKDHB* on chromosomal 6, which resulted in a C -terminal truncated form of the protein. B) RNA-seq confirmed the WGS data and showed a dominant allelic expression of the corresponding fusion transcript, with exons 1-17 of *EP300* fused to a short stretch of intronic sequence of *BCKDHB*. Red or purple indicated orientations of reads (forward or reversed).

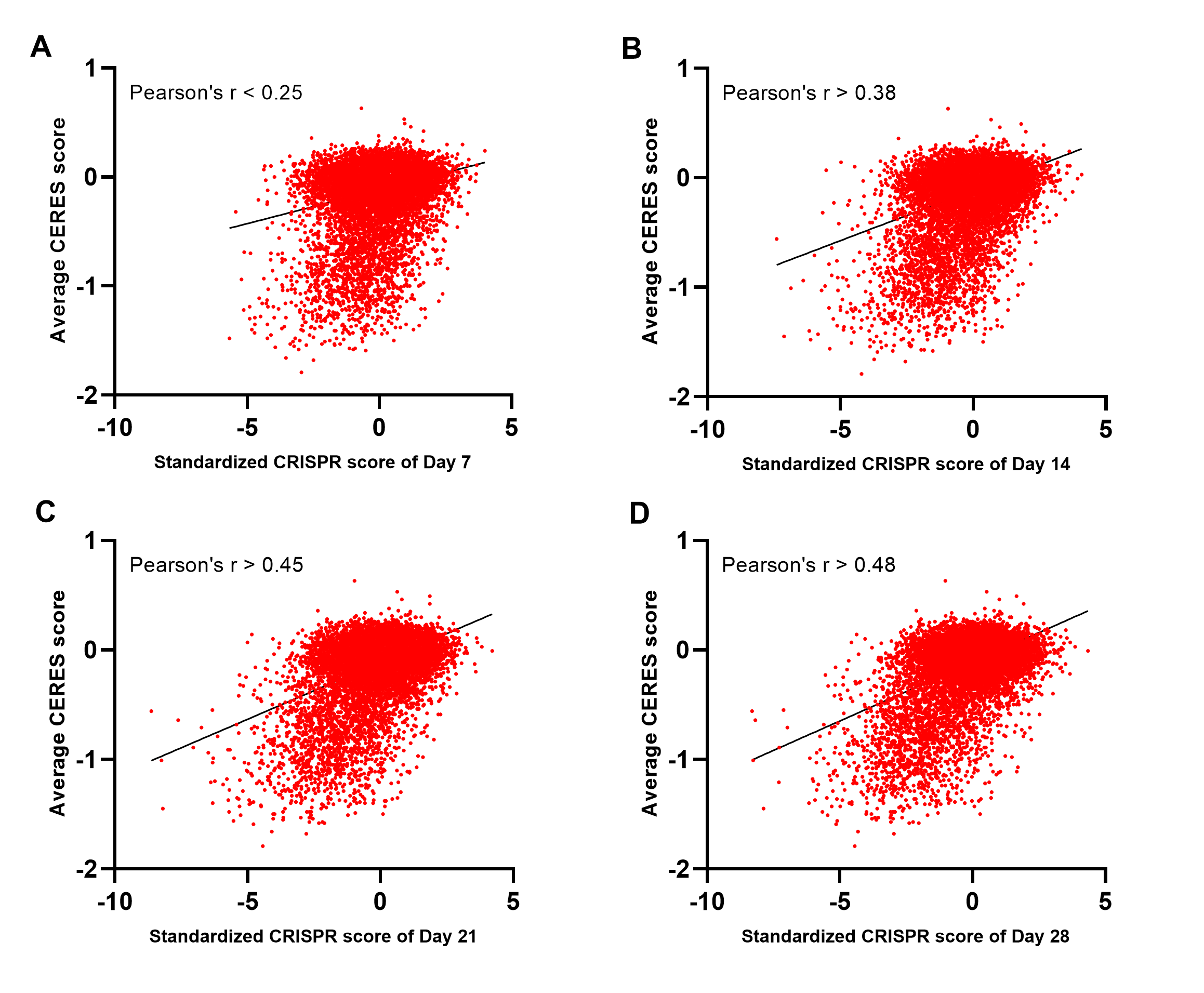

**Figure S3 | Correlation between the the standardized CRISPR scores of RC-K8 and the average CERES scores.** Scatter plots showing the correlation between the standardized CRISPR scores of RC-K8 at different time points and the average CERES scores (4). Pearson correlation values are indicated.

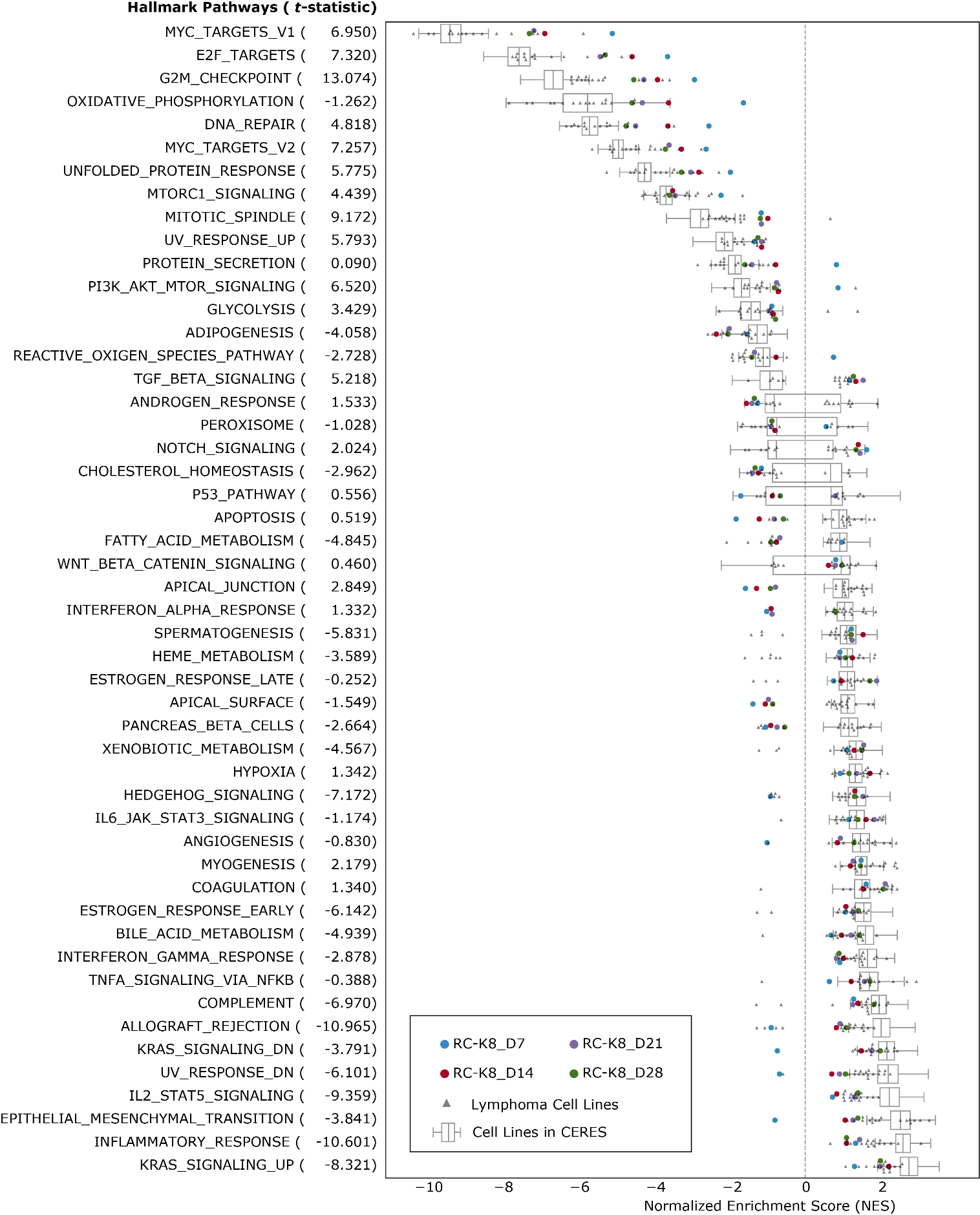

**Figure S4 | Gene set enrichment analysis (GSEA) of 50 hallmark pathways**. Normalized gene enrichment scores based on RC-K8 screening results at four time points (coloured dots), compared to enrichment scores based on previously reported screens from B-cell lymphoma cell lines (grey triangles) (5,6) and 341 cancer cell lines from the CERES data (box-plot) (4). Pathways are sorted by median enrichment scores of all cell lines. T-statistics were calculated by comparing the enrichment scores between lymphoma cell lines and CERES cell lines, indicating the specificity of enrichments towards B-cell lymphoma.

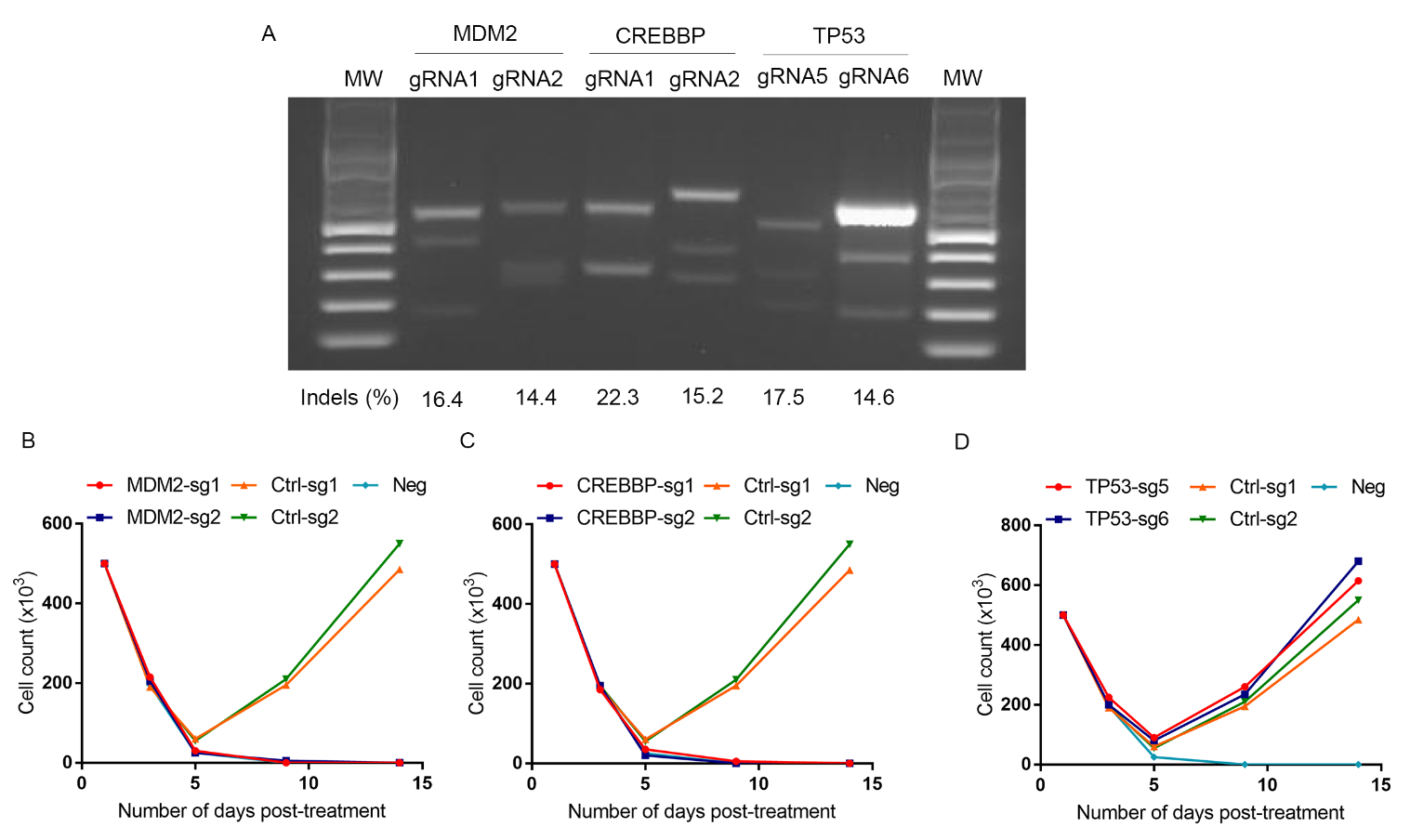

**Figure S5| Validation of top candidates.** (**A**) SURVEYOR assay was used to determine the indel ratio of transfected RC-K8 cells. Cas9-mediated cleavage efficiency (indel ratio) was calculated on the basis of integrated intensity of gel binds. Genomic DNA was extracted from cells on day 5 post-transduction. (**B-D**) RC-K8 cells transduced with *MDM2*-targeting lentiCRISPRs (**B**) and *CREBBP*-targeting lentiCRISPRs (**C**) showed a significant growth inhibition, whereas cells transduced with lentiCRISPR vectors did not. (**D**) RC-K8 cells transduced with *TP53*-targeting lentiCRISPRs showed no changes in growth compared to cells transduced with lentiCRISPR vectors.

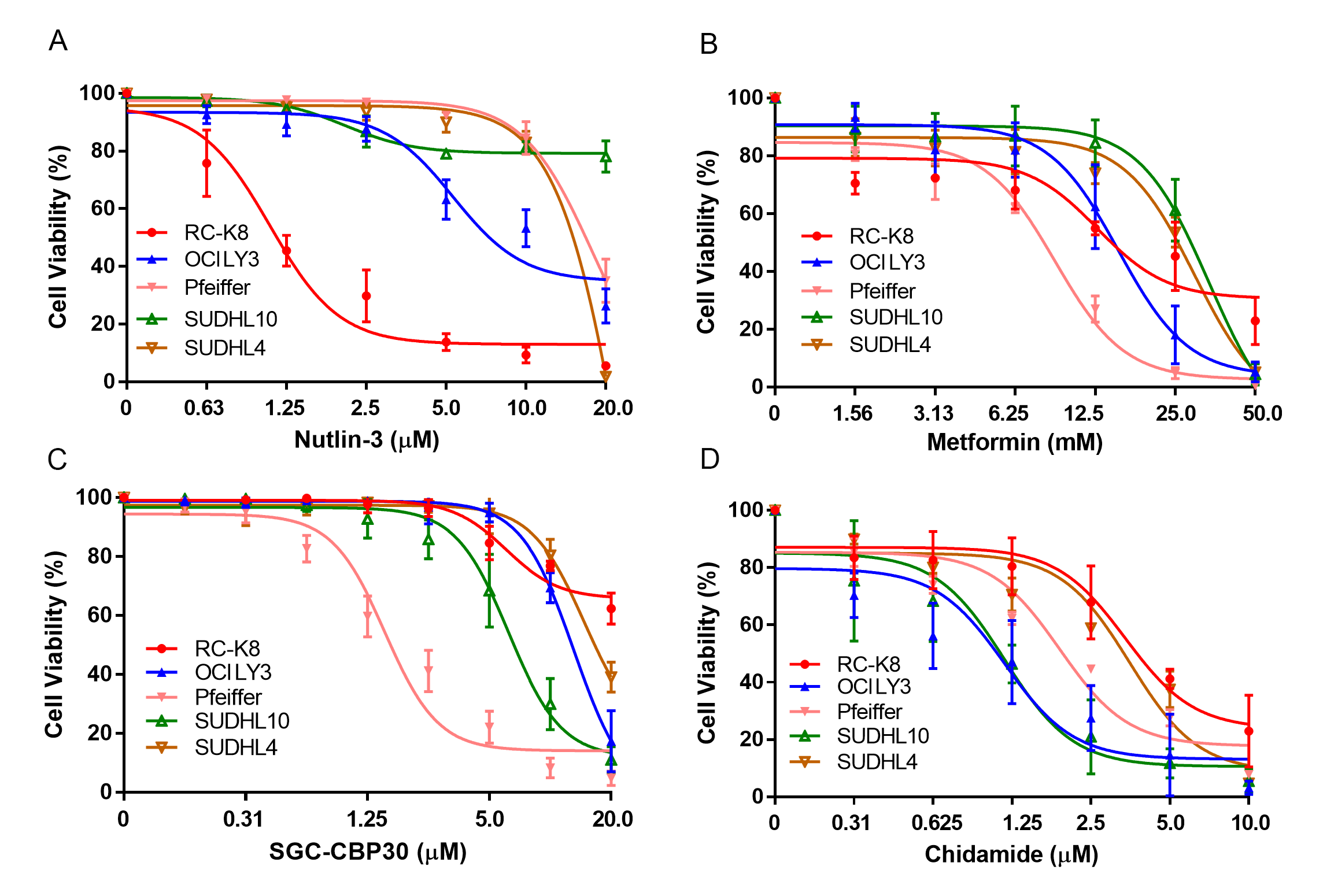

**Figure S6 | Cell viability tests in DLBCL cell lines**. Cells were treated with indicated inhibitors for 72h. A) MDM2 inhibition by small molecule nutlin-3; B) OXPHOS inhibition by biguanide metformin; C) BRD-domain inhibition by small molecule SGC-CBP30; D) Histone deacetylase (HDAC) inhibition by Chidamide. The data represent the means ± SD of three independent experiments.

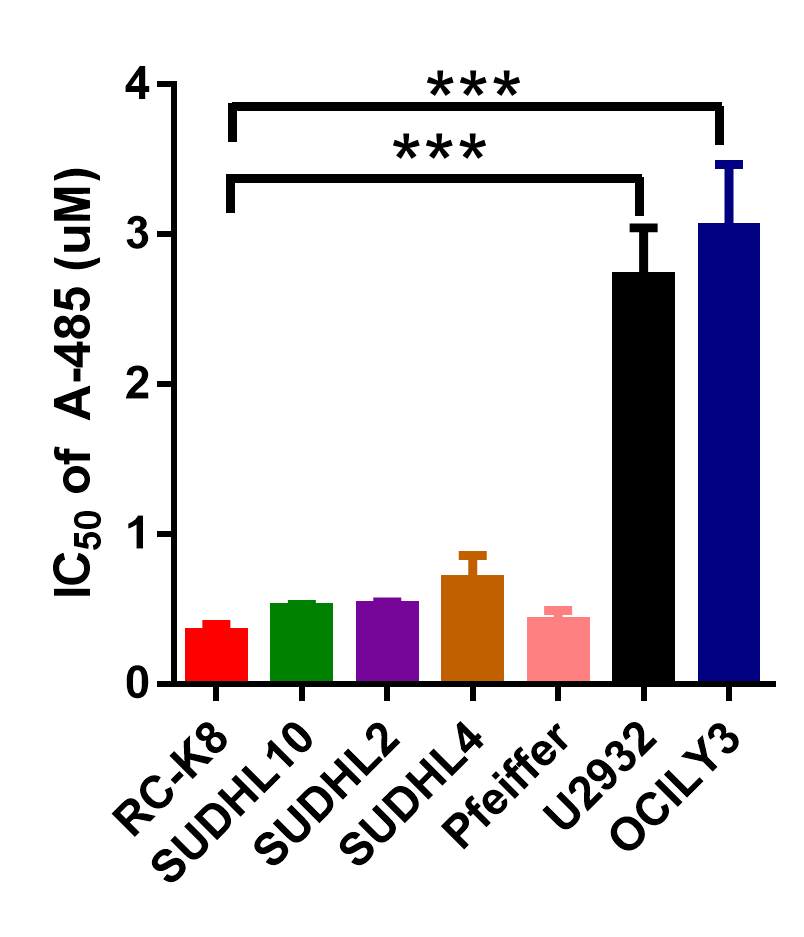

**Figure S7 | The IC50 value of A-485 in different DLBCL cell lines.** Compared to cell lines (OCILY3 and U2932) with wild-type HATs, the five HAT-mutant cell lines displayed higher sensitivity to the inhibition, with RC-K8 cells exhibited the lowest IC50 value. ***P < 0.001, one-way ANOVA with Tukey. The data represent the means ± SD of three independent experiments.

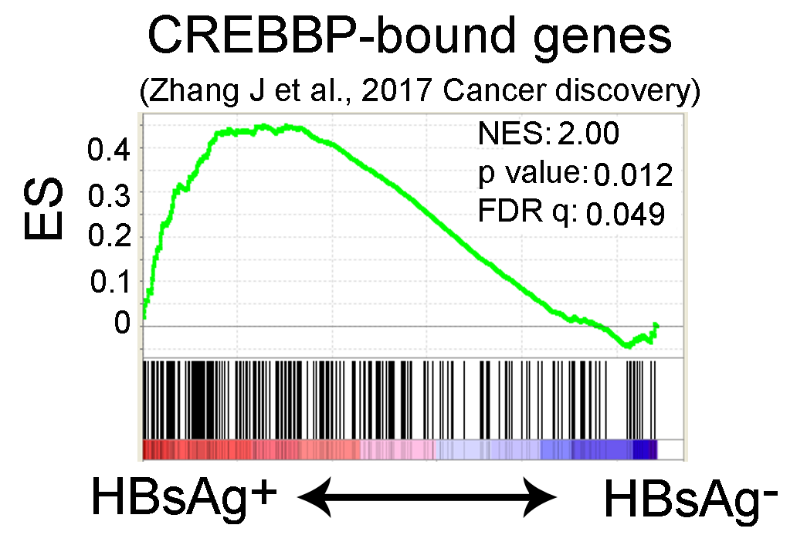

**Figure S8 | CREBBP-bound gene set was significantly upregulated in HBsAg^+^ DLBCLs.** GSEA plot illustrating the enrichment of CREBBP-bound genes in HBsAg^+^ DLBCLs compared to HBsAg^-^ DLBCLs. RNAseq data was described previously (7) and re-analyzed here. GSEA were performed with 1000 sample permutations. Enrichments were considered significant if FDR q < 0.25. ES, enrichment score. NES: normalized enrichment score.

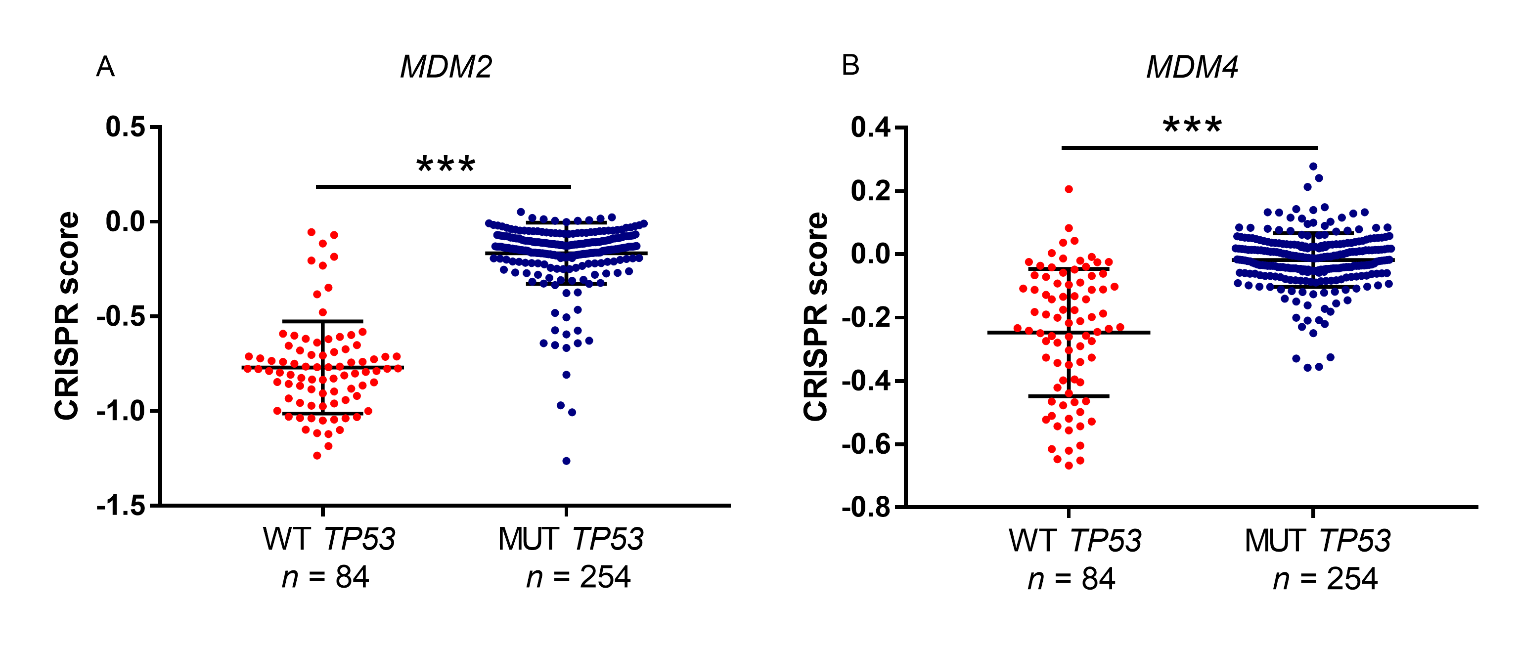

**Figure S9| *MDM2* and *MDM4* were more essential in cancer cell lines with WT *TP53*.** In CERES dataset (4), CRISPR score of *MDM2* (A) and *MDM4* (B) were significantly lower in cancer cell lines with WT *TP53* (*n* = 84) than MUT *TP53* (*n* = 254). Student’s *t* test, ****P* < 0.001.

**References:**

1.	Chen X, Schulz-Trieglaff O, Shaw R, Barnes B, Schlesinger F, Kallberg M*, et al.* Manta: rapid detection of structural variants and indels for germline and cancer sequencing applications. Bioinformatics **2016**;32:1220-2

2.	Garbati MR, Thompson RC, Haery L, Gilmore TD. A rearranged EP300 gene in the human B-cell lymphoma cell line RC-K8 encodes a disabled transcriptional co-activator that contributes to cell growth and oncogenicity. Cancer Lett **2011**;302:76-83

3.	Garbati MR, Alco G, Gilmore TD. Histone acetyltransferase p300 is a coactivator for transcription factor REL and is C-terminally truncated in the human diffuse large B-cell lymphoma cell line RC-K8. Cancer Lett **2010**;291:237-45

4.	Meyers RM, Bryan JG, McFarland JM, Weir BA, Sizemore AE, Xu H*, et al.* Computational correction of copy number effect improves specificity of CRISPR-Cas9 essentiality screens in cancer cells. Nat Genet **2017**;49:1779-84

5.	Phelan JD, Young RM, Webster DE, Roulland S, Wright GW, Kasbekar M*, et al.* A multiprotein supercomplex controlling oncogenic signalling in lymphoma. Nature **2018**;560:387-91

6.	Reddy A, Zhang J, Davis NS, Moffitt AB, Love CL, Waldrop A*, et al.* Genetic and functional drivers of diffuse large B cell lymphoma. Cell **2017**;171:481-94 e15

7.	Ren W, Ye X, Su H, Li W, Liu D, Pirmoradian M*, et al.* Genetic landscape of hepatitis B virus-associated diffuse large B-cell lymphoma. Blood **2018**;131:2670-81
